# A deep learning framework for predicting human essential genes from population and functional genomic data

**DOI:** 10.1101/2021.12.21.473690

**Authors:** Troy M. LaPolice, Yi-Fei Huang

## Abstract

Being able to predict essential genes intolerant to loss-of-function (LOF) mutations can dramatically improve our ability to identify genes associated with genetic disorders. Numerous computational methods have recently been developed to predict human essential genes from population genomic data; however, the existing methods have limited power in pinpointing short essential genes due to the sparsity of polymorphisms in the human genome. Here we present an evolution-based deep learning model, DeepLOF, which integrates population and functional genomic data to improve gene essentiality prediction. Compared to previous methods, DeepLOF shows unmatched performance in predicting ClinGen haploinsufficient genes, mouse essential genes, and essential genes in human cell lines. Furthermore, DeepLOF discovers 109 potentially essential genes that are too short to be identified by previous methods. Altogether, DeepLOF is a powerful computational method to aid in the discovery of essential genes.

## Introduction

Loss-of-function (LOF) mutations, including stop-gain, splice-site, and frameshift mutations, play a key role in the etiology of genetic disorders. While it is relatively straightforward to identify LOF mutations in protein-coding genes, it is challenging to infer their effects on evolutionary fitness and disease risk. Several computational methods^1–7^ have recently been developed to predict human essential genes based on the premise that LOF mutations causing early-onset disorders may be subject to negative selection in human populations^8,9^. Based on large-scale population genomic data, such as gnomAD^5^, these methods seek to identify LOF-intolerant genes where the observed number of LOF variants is significantly smaller than the expected number under a neutral mutation model. It has been shown that LOF-intolerant genes predicted by these methods are enriched with haploinsufficient genes associated with Mendelian disorders^1–7^. Furthermore, *de novo* LOF mutations in probands with autism^10,11^, schizophrenia^12,13^, and severe developmental disorders14 are significantly overrepresented in LOF-intolerant genes. Therefore, population genetics-based prediction of LOF-intolerant genes is a powerful strategy to discover haploinsufficient genes associated with human disease.

Despite the recent success of population genetics-based gene essentiality prediction, the statistical power of existing methods may heavily depend on the length of a gene^5,9,15^. Specifically, a long gene typically has a large expected number of LOF variants under a neutral mutation model. Thus, when we compare the observed number of LOF variants with the expected one, it is relatively easy to reject the null hypothesis of neutral evolution in a long gene. In contrast, a short gene is expected to have only a handful of LOF variants. Therefore, in a short gene it is difficult to distinguish the depletion of LOF mutations caused by negative selection from that by chance alone, which may hinder the discovery of many essential genes of short length in the human genome.

Complementary to population genomic data that manifest natural selection at the organism level, functional genomic assays, such as RNA-seq, ChIP-seq, and genome editing, provide rich information on the molecular functions of protein-coding genes. Thus, functional genomic data may also be utilized to predict gene essentiality. Based on this idea, several supervised methods have been developed to predict essential genes from functional genomic features^15–20^. Unlike population genetics-based methods, the predictive power of genomic feature-based methods may not heavily depend on the length of a gene. However, because functional genomic data are often from cell lines, gene scores solely derived from functional genomic features may not always be indicative of gene essentiality at the organism level.

In the current study, we propose that integrating population and functional genomic data may improve gene essentiality prediction. To this end, we introduce DeepLOF, an evolution-based deep learning model for predicting human genes intolerant to LOF mutations. By combining a deep neural network and a population genetics-based likelihood function, DeepLOF can integrate genomic features and population genomic data to predict LOF-intolerant genes without human-labeled training data. Compared to previous methods, DeepLOF shows unmatched performance in predicting ClinGen haploinsufficient genes^21^, human orthologs of mouse essential genes^22^, and genes essential to the survival of cell lines^23^. Furthermore, using DeepLOF we identify 109 LOF-intolerant genes of short length missed by previous methods. The 109 novel LOF-intolerant genes are enriched with essential genes and are depleted in benign genomic deletions. Taken together, DeepLOF is a powerful deep learning framework to predict essential genes in the human genome.

## Results

### Overview of the DeepLOF model

DeepLOF is an evolution-based deep learning model for inferring protein-coding genes intolerant to LOF mutations. The key variable of interest in DeepLOF is *η, i.e.,* the relative rate of LOF variants in a gene with respect to the expected number of LOF variants under a neutral mutation model. A smaller *η* indicates that a gene has a lower rate of LOF variants after adjusting for neutral evolutionary factors, such as mutation rate and genetic drift. Thus, a smaller *η* indicates stronger negative selection against LOF variants. To take into account the uncertainty of *η*, DeepLOF treats *η* as a random variable at the gene level. To integrate genomic features and population genomic data in a Bayesian manner, DeepLOF combines a feedforward neural network and a population genetics-based likelihood function (Fig. 1).

**Figure 1:**
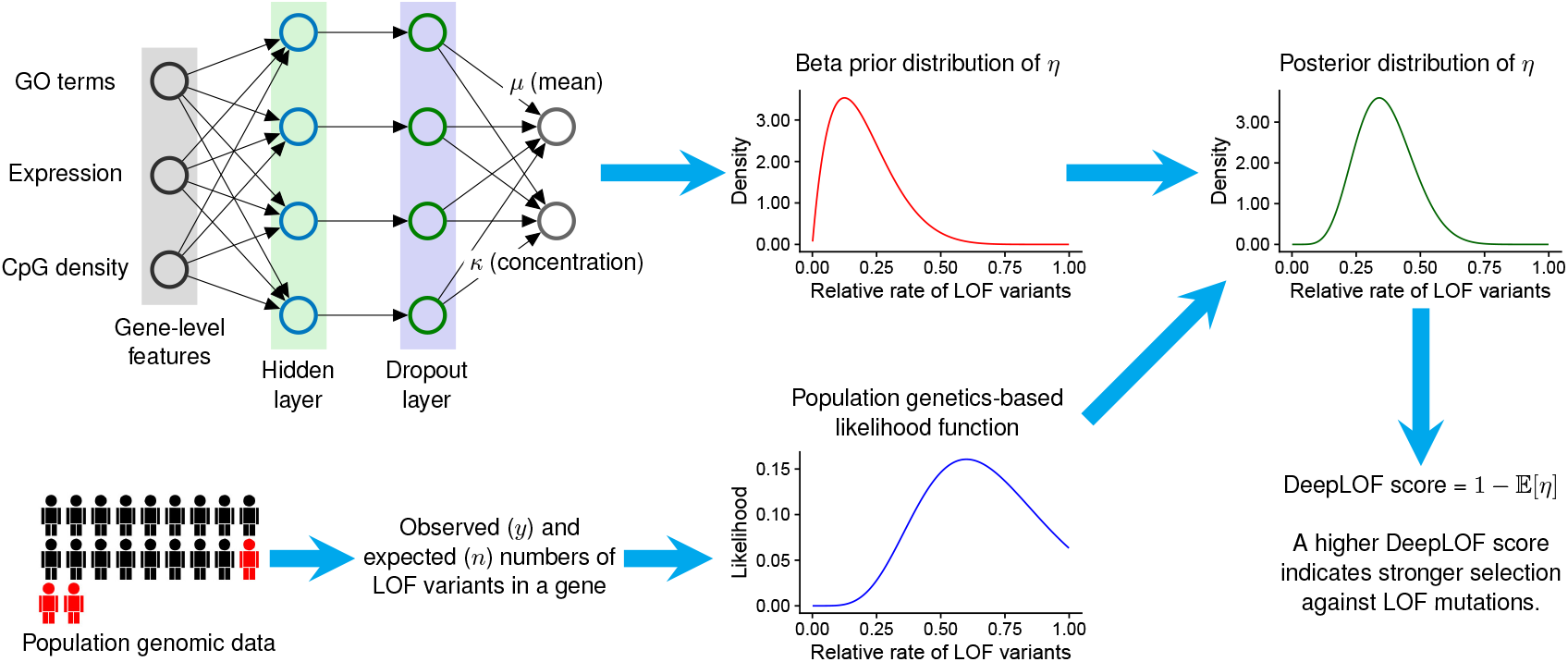
Overview of the DeepLOF model. DeepLOF combines a feedforward neural network and a population genetics-based likelihood function to infer the relative rate of LOF variants in a gene (*η*) with respect to the expected number under a neutral mutation model (*n*). The feedforward neural network transforms genomic features into a beta prior distribution of *η*, which represents our belief about *η* based on genomic features. The population genetics-based likelihood function describes the probability of observing *y* LOF variants in a gene conditional on *η* and *n*, which represents our belief about *η* based on population genomic data. Finally, DeeLOF combines the prior distribution and the likelihood function to compute the posterior distribution of *η*. The DeepLOF score is defined as 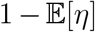, where 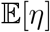 is the mean of *η* under the posterior distribution.

In this hybrid framework, the feedforward neural network consists of a sequence of neural network layers, which together transform genomic features into the beta prior distribution of *η* (Fig. 1). The genomic features include gene ontology (GO) terms^24^, epigenomic data, gene expression patterns, and several other gene-level features potentially predictive of LOF intolerance. The outputs of the feedforward neural network are the mean and concentration parameters of the beta distribution, which represents our belief about *η* based on genomic features. In addition, the population genetics-based likelihood function describes the probability of observing *y* LOF variants in a gene given *η* and n, where *n* is the expected number of LOF variants in the same gene under a neutral mutation model (Fig. 1). Thus, the likelihood function represents evidence for LOF intolerance based on population genomic data.

Using Bayes’ rule, DeepLOF combines the neural network-based beta prior distribution with the population genetics-based likelihood function to obtain a posterior distribution of *η*, which represents our belief about LOF intolerance after integrating genomic features and population genomic data. Denoting 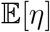 as the expectation of *η* under the posterior distribution, we define the DeepLOF score as 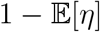, which can be interpreted as the proportion of LOF mutations purged by negative selection in a gene. Thus, a higher DeepLOF score indicates a higher level of LOF intolerance. We estimate model parameters, including the weights and biases of the feedforward neural network, using stochastic gradient descent on a loss function that integrates the feedforward neural network and the likelihood function.

### DeepLOF elucidates genomic features predictive of LOF-intolerant genes

We trained the DeepLOF model on the observed and expected numbers of LOF variants in 19,197 human genes and 18 genomic features (Supplementary Data 1). The observed number of LOF variants in each gene was from the exomes of 125,748 healthy individuals in the gnomAD database^5^. The expected number of LOF variants in each gene was from a neutral mutation model developed by gnomAD^5^, which took into account the impact of trinucleotide sequence context, CpG methylation level, local mutation rate, and site-wise sequencing coverage on the occurrence of variants. The 18 genomic features included five epigenomic features^20^, four gene categories associated with developmental processes^25^, three protein annotations^26–28^, two phastCons conservation scores^15,29^, two gene expression features^30,31^, the promoter CpG density^15^, and the UNEECON-G score^32^. A detailed description of these genomic features is available in Supplementary Table 1. We used 80% randomly selected genes as a training set and used the remaining 20% genes as a validation set for hyperparameter tuning.

To obtain insights into which genomic features may be predictive of gene-level intolerance to LOF mutations, we trained a linear DeepLOF model without hidden layer in the feedforward neural network. While the linear DeepLOF model may not provide most accurate predictions of LOF intolerance, it allows us to estimate the association of each genomic feature with LOF intolerance. Specifically, in the linear DeepLOF model, we defined the contribution score of a genomic feature as the negative value of its weight with respective to the mean of the beta prior distribution of η. The absolute value of a contribution score indicates the strength of association between a feature and LOF intolerance, whereas the sign of the contribution score indicates the direction of association.

Among the 18 genomic features, the UNEECON-G score had the strongest positive association with LOF intolerance (Fig. 2a). Because the UNEECON-G score is a measure of a gene’s intolerance to missense mutations, it corroborates a previous finding that missense intolerance is strongly correlated with LOF intolerance at the gene level^32^. Two GO categories^24^, *i.e.*, central nervous system development and embryo development, and the Reactome category of nervous system development33 had strong positive associations with LOF intolerance, suggesting that developmental genes are highly intolerant to LOF mutations. Two protein annotations, i.e., transcription factor26 and protein complex^27^, also had strong positive associations with LOF intolerance, suggesting that genes encoding transcription factors or subunits of protein complexes may be more intolerant to LOF mutations than other protein-coding genes. In agreement with previous studies^15,20^, epigenomic features in a gene’s promoter, including the signals of H3K9ac, H3K27me3, and H3K4me3 histone modifications^20^, and the promoter CpG density15 had positive associations with LOF intolerance. Furthermore, the phastCons score29 in a gene’s promoter had a positive association with LOF intolerance, suggesting that genes with conserved promoter sequences may be intolerant to LOF mutations. Finally, tissue specificity34 had a negative association with LOF intolerance, suggesting that housekeeping genes may be more intolerant to LOF mutations than tissue-specific genes.

**Figure 2:**
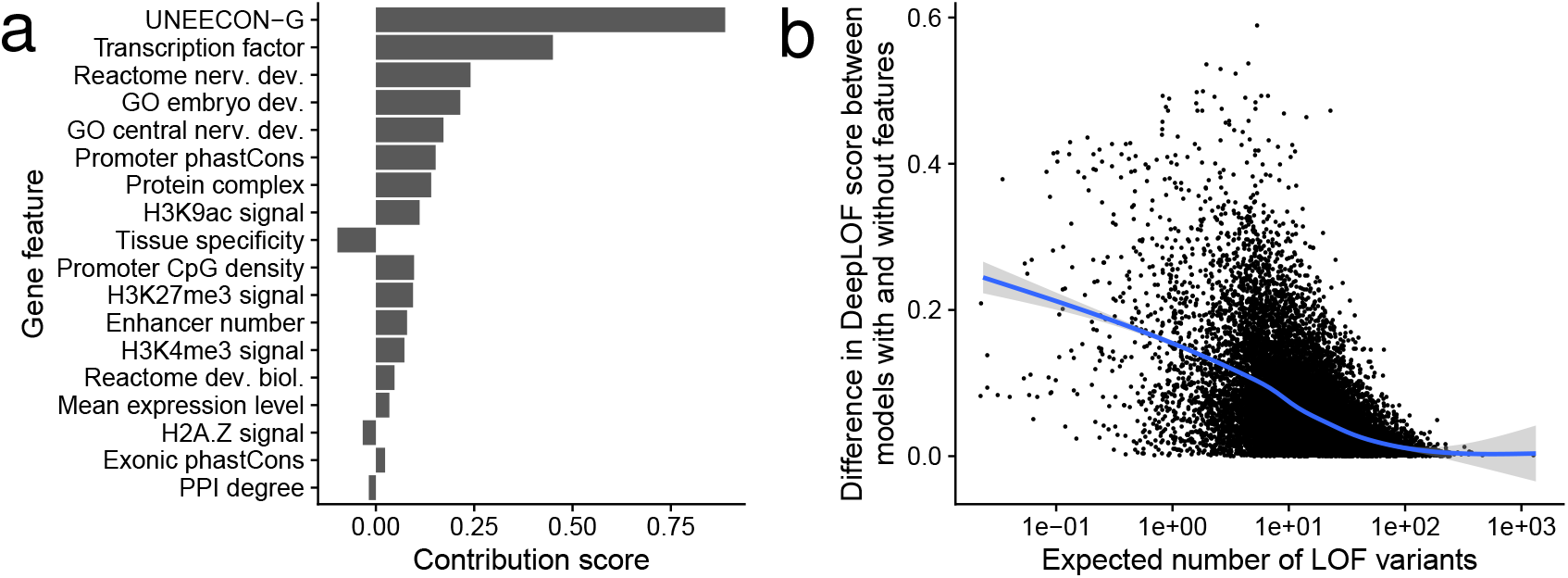
Impact of genomic features on the inference of LOF intolerance. (a) Association of genomic features with LOF intolerance. We define the contribution score of a genomic feature as the negative weight of the feature in the linear DeepLOF model. The absolute value of a contribution score indicates the strength of association between a feature and LOF intolerance, whereas the sign of a contribution score indicates the direction of association. (b) DeepLOF automatically adjusts the relative importance of genomic features in a gene length-dependent manner. The *x* axis represents the expected number of LOF variants. The *y* axis represents the absolute difference in DeeLOF score between the linear DeepLOF model with genomic features and the counterpart model without genomic features. A higher absolute difference in DeepLOF score indicates a stronger impact of genomic features on the inference of LOF intolerance. Each dot represents a gene. The blue and grey curves represent the fit of the generalized additive model with integrated smoothness and its 95% confidence interval^58^.

### DeepLOF automatically adjusts the relative importance of genomic features and population genomic data in a gene length-dependent manner

Because DeepLOF uses Bayes’ rule to infer the distribution of *η*, we hypothesized that, similar to other Bayesian models, DeepLOF may automatically adjust the relative importance of the beta prior distribution in a data-dependent manner. Specifically, in a long gene where a large number of LOF variants is expected under a neutral mutation model, the posterior distribution of *η* may be dominated by the the population genetics-based likelihood function. Thus, DeepLOF may primarily leverage population genomic data to predict long LOF-intolerant genes. Because population genomic data are indicative of negative selection at the organism level, this may allow DeepLOF to unbiasedly infer LOF intolerance at the organism level in long genes. Conversely, in a short gene where a small number of LOF variants is expected under a neutral mutation model, the posterior distribution of *η* may be dominated by the beta prior distribution of η. Thus, DeepLOF may automatically upweight genomic features to improve the inference of LOF intolerance in short genes.

To test this hypothesis, we retrained the linear DeepLOF model without using any genomic features. This model effectively assumed an identical prior distribution of *η* across genes and solely used population genomic data to infer LOF intolerance. We computed the absolute difference in DeepLOF score between the linear DeepLOF model with genomic features and the model without genomic features, which indicates the relative importance of genomic features in the inference of LOF intolerance. We observed that the absolute difference in DeepLOF score was negatively correlated with the expected number of LOF variants in a gene (Fig. 2b), supporting our hypothesis that DeepLOF automatically upweights genomic features in short genes to improve the prediction of LOF intolerance.

### DeepLOF shows unmatched performance in predicting essential genes intolerant to LOF mutations

We hypothesized that, by integrating a large number of genomic features and population genomic data, DeepLOF may show improved performance in predicting essential genes. To test this hypothesis, we obtained three sets of essential genes, including 311 ClinGen haploinsufficient genes^21^, 397 human orthologs of mouse essential genes where heterozygous knockouts resulted in lethality^22^, and 683 human genes essential to the survival of cell lines^23^. For each essential gene set, we constructed a nonessential gene set of matching size. To this end, we used MatchIt35 to match each essential gene with a putatively nonessential gene of similar number of LOF variants.

We trained a nonlinear DeepLOF model with hidden layer and observed that the nonlinear DeepLOF model had a lower loss than the linear DeepLOF model in the validation set. Thus, we used scores from the nonlinear DeepLOF model in downstream analysis (Supplementary Data 2). We compared the performance of DeepLOF with eight alternative methods in distinguishing essential genes from matched nonessential genes. The eight alternative methods included two measures of gene-level intolerance to LOF mutations (LOEUF5 and pLI2), three measures of gene-level intolerance to missense mutations (mis-z^1^, GeVIR^6^, and UNEECON-G^32^), and three metrics that considered both LOF intolerance and missense intolerance (RVIS^36^, VIRLOF^6^, CoNeS^7^). In the prediction of ClinGen haploinsufficient genes, DeepLOF showed substantially better performance than the other methods as evidenced by its significantly higher area under the receiver operating characteristic curve (AUC) (Fig. 3a; Supplementary Table 2). In particular, DeepLOF showed unmatched performance when the false positive rate was low. For instance, the true positive rate of DeepLOF was approximately 0.5 at a false positive rate of 0.05 (Fig. 3b), which was about 50% higher than that of the second best method (LOEUF; true positive rate = 0.33). DeepLOF also outperformed the other methods in predicting human orthologs of mouse essential genes and human genes essential for the survival of cell lines (Fig. 3c & d; Supplementary Table 2). In sum, DeepLOF had superior performance in predicting essential genes.

**Figure 3:**
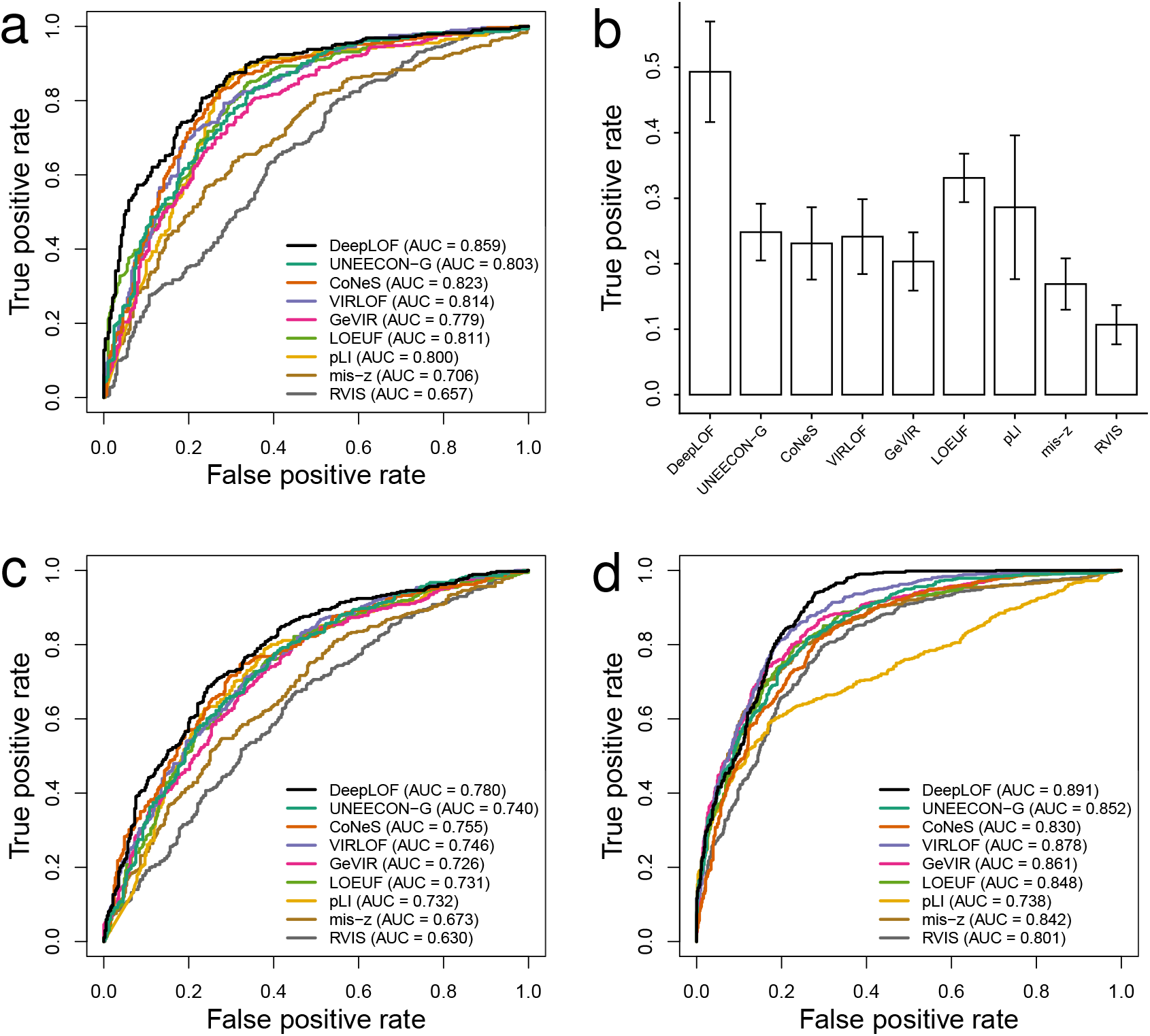
Predictive power of DeepLOF and alternative methods in distinguishing essential genes from putatively nonessential genes. (a) Performance in predicting ClinGen haploinsufficient genes. (b) True positive rates in predicting ClinGen haploinsufficient genes at a fixed false negative rate of 5%. Error bars represent bootstrap standard errors of true positive rates. (c) Performance in predicting human orthologs of mouse essential genes. (d) Performance in predicting human genes essential for the survival of cell lines.

Because the DeepLOF score is a measure of LOF intolerance, we hypothesized that it might not be the best method in predicting disease genes via a mechanism different from haploinsufficiency. To test this hypothesis, we obtained 364 OMIM dominant-negative genes where a heterozygous mutation may adversely affect the function of the wild-type allele in the same individual through interlocus or intralocus interactions^36–38^. In agreement with our hypothesis, UNEECON-G instead of DeepLOF showed the best performance in predicting dominant-negative genes (Supplementary Fig. 1), suggesting that missense intolerance scores, such as UNEECON-G, might be better predictors of dominant-negative genes than LOF intolerance scores.

### DeepLOF predicts 109 novel LOF-intolerant genes of short length

Previous predictions of LOF intolerance are often biased towards longer genes because it is easier to reject neutral evolution when the expected number of LOF variants is high6'^15^. In contrast, by leveraging a genomic feature-based prior distribution, DeepLOF may have higher power to predict LOF-intolerant genes of short length. To test this hypothesis, we examined LOF-intolerant genes predicted by four methods, including DeepLOF, CoNeS, LOEUF, and VIRLOP, which showed better performance than other methods in predicting ClinGen haploinsufficient genes (Fig. 3a). Using a previously established cutoff of 0.35 (ref. 5), we obtained 2,835 LOF-intolerant genes from LOEUF. Using comparable cutoffs, we obtained 2,817, 2,847, and 2,817 LOF-intolerant genes from DeepLOF, CoNeS, and VIRLOP, respectively. Because previous studies suggested that the difficulty of LOF intolerance prediction mainly occurred in genes with ≤ 10 expected LOF variants^5,15^, we focused on predicted LOF-intolerant genes with ≤ 10 expected LOF variants in downstream analysis.

In total, 452 genes with ≤ 10 expected LOF variants were predicted to be LOF-intolerant by at least one method (Supplementary Data 3). DeepLOF predicted that 364 genes with ≤ 10 expected LOF variants were LOF-intolerant, which was the largest number among all the four methods (Fig. 4a). Also, 109 LOF-intolerant genes, or 24.1% of the total number (109/452), were uniquely predicted by DeepLOF (Fig. 4a). Because we used comparable cutoffs for the four methods, these results suggest that DeepLOF may have much higher power to pinpoint LOF-intolerant genes of short length.

**Figure 4:**
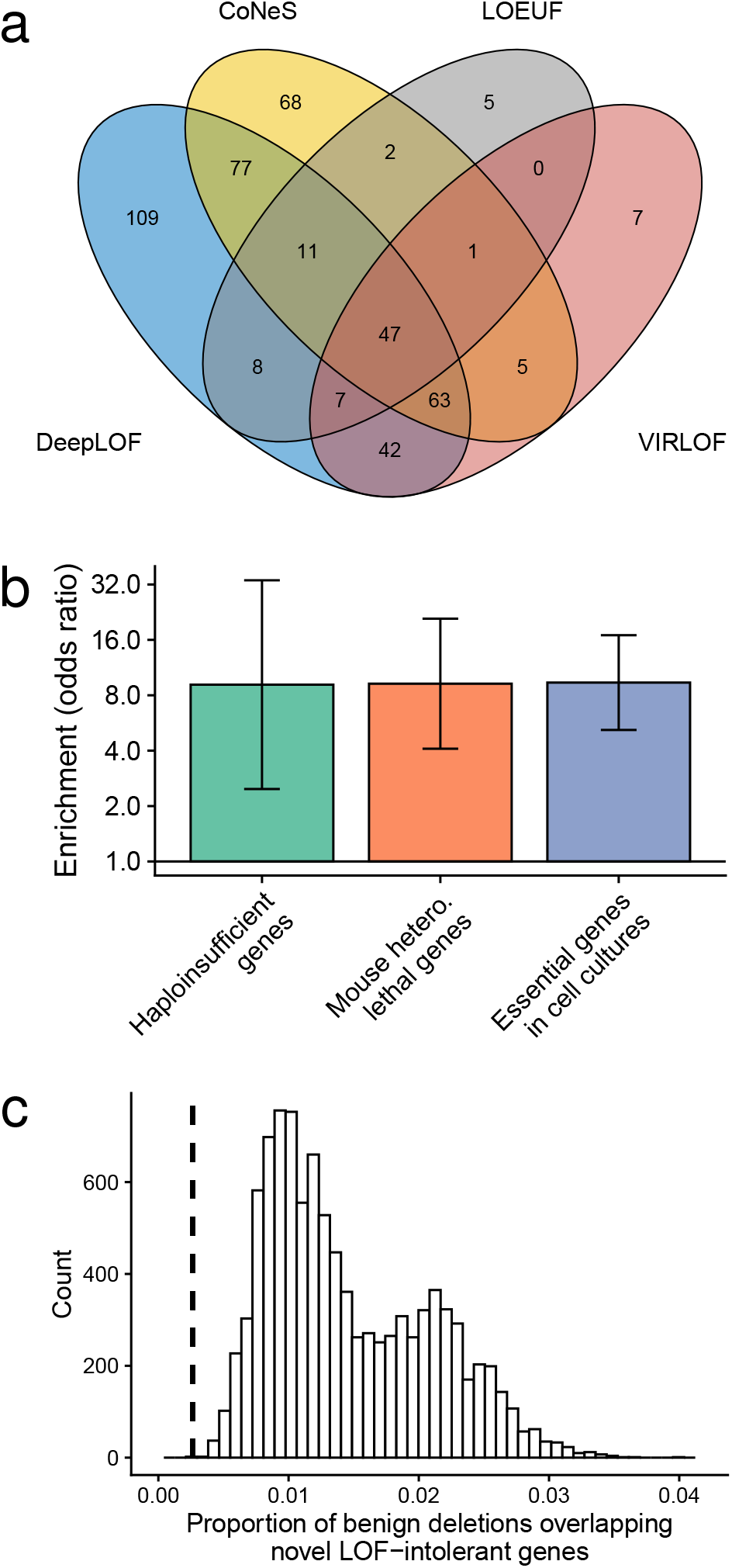
Predicted LOF-intolerant genes with ≤ 10 expected LOF variants. (a) Venn diagram of predicted LOF-intolerant genes. (b) Enrichment of essential genes in the 109 LOF-intolerant genes uniquely predicted by DeepLOF. Error bars represent 95% confidence intervals. (c) Proportion of benign genomic deletions overlapping the 109 LOF-intolerant genes uniquely predicted by DeepLOF. The histogram represents the null distribution of the proportion from a permutation test, and the dashed vertical line represents the observed proportion in empirical data.

We examined the enrichment of ClinGen haploinsufficient genes, human orthologs of mouse essential genes, and human genes essential for the survival of cell lines in the 109 novel LOF-intolerant genes predicted by DeepLOF, using genes with ≤ 10 expected LOF variants but not predicted to be LOF-intolerant by any method as a background set. We observed that the 109 novel LOF-intolerant genes were significantly enriched with essential genes (Fig. 4b), highlighting that the novel LOF-intolerant genes predicted by DeepLOF may play key roles in important biological processes.

Finally, we hypothesized that the 109 novel LOF-intolerant genes predicted by DeepLOF might be depleted in benign genomic deletions due to the detrimental effects of deletions overlapping LOF-intolerant genes. To test this hypothesis, we obtained 5,649 benign genomic deletions overlapping protein-coding genes from dbVar^39^. We observed that 0.27% of benign deletions overlapped the 109 novel LOF-intolerant genes (Fig. 4c). To evaluate whether the proportion of overlapping was smaller than the expectation under a null model that postulated the 109 genes to be nonessential, we performed a permutation test with 10,000 permutations. In each permutation, we randomly selected 109 genes with ≤ 10 LOF variants and computed the proportion of benign genomic deletions overlapping the random genes. The mean proportion of benign deletions overlapping random genes was 1.46% (Fig. 4c), which was 5.4 fold higher than the observed proportion in empirical data (1.46% vs. 0.27%; *P* = 0; one-tailed permutation test). Thus, benign genomic deletions were depleted with the 109 novel LOF-intolerant genes predicted by DeepLOF.

## Discussion

In the current work, we present an evolution-based machine learning framework, DeepLOF, for predicting human genes intolerant to LOF mutations. Unlike previous LOF intolerance scores, such as pLI and LOEUF, the DeepLOF model leverages both population and functional genomic data to predict LOF intolerance. Therefore, DeepLOF may be particularly powerful in predicting short essential genes without sufficient polymorphisms for selection inference. Furthermore, unlike supervised methods, DeepLOF does not use known essential genes as training data and, thus, may not suffer from label leakage and other pitfalls of supervised machine learning^40^.

The linear DeepLOF model without hidden layer allows us to directly estimate the association of a genomic feature with LOF intolerance after adjusting for other genomic features (Fig. 2a). Using this approach, we show that the UNEECON-G score has the strongest positive association with LOF intolerance, which suggests that missense intolerance scores may also be informative of gene-level intolerance to LOF mutations. Because there are typically more missense variants than LOF variants in a gene under a neutral mutation model, the sample size for missense intolerance inference is larger than that for LOF intolerance inference. Therefore, it may be easier to reliably estimate missense intolerance than LOF intolerance, and in turn it may be beneficial to incorporate missense intolerance scores, such as UNEECON-G, into computational pipelines for LOF variant interpretation.

We also show that genes encoding transcription factors or protein complex subunits and genes associated with developmental processes may be highly intolerant to LOF mutations (Fig. 2a). Previous studies have shown that many transcription factors are haploinsufficient and are associated with dominant genetic disorders^41^. Transcription factors often cooperatively bind to regulatory sequences, which may result in a sigmoid-shaped dose-response curve^42,43^. Therefore, transcription factors may be particularly susceptible to heterozygous knockouts. Also, in agreement with our observation, it has been shown that many protein complex subunits are haploinsufficient44 because the reduced expression of a subunit may lead to a stoichiometric imbalance between different subunits of the same protein complex^43^. Finally, in agreement with our observation, it has been found that many developmental genes are haploinsufficient^45^, highlighting that developmental processes may be particularly sensitive to reduced gene dosage.

While both genomic features and population genomic data are predictive of LOF intolerance, their relative importance may depend on the length of a gene. In short genes where population genomic data provide limited information on negative selection, it is critical to incorporate genomic features to improve the inference of LOF intolerance. In contrast, in long genes where polymorphisms are abundant, population genomic data may be more informative than genomic features because they directly reflect LOF intolerance at the organism level. Because DeepLOF infers the relative rate of LOF variants, *η*, in a Bayesian manner, it can automatically adjust the relative importance of genomic features and population genomic data to optimize LOF intolerance inference in a data-dependent manner (Fig. 2b).

By integrating genomic features and population genomic data, DeepLOF outperforms alternative methods in predicting essential genes (Fig. 3). Because most variants in the gnomAD database are of low allele frequency^5^, the DeepLOF score may be indicative of negative selection against LOF variants in their heterozygous state. Thus, it shows unmatched performance in predicting ClinGen haploinsufficient genes. In contrast, DeepLOF may not be a powerful method to predict dominant-negative disease genes (Supplementary Fig. 1), highlighting that it is critical to take into account the genetic mechanism of disease in gene prioritization^46^.

Because DeepLOF leverages genomic features to improve the inference of LOF intolerance in short genes, DeepLOF has predicted the largest number of short LOF-intolerant genes compared to other methods (Fig. 4a). Furthermore, DeepLOF has predicted 109 novel LOF-intolerant genes of short length. These novel LOF-intolerant genes are enriched with essential genes and are depleted in benign genomic deletions (Fig. 4b & c), implicating that they may play an underappreciated role in human disease.

## Methods

### Details of the DeepLOF model

We denote *η_i_* as the relative rate of observed LOF variants in gene *i* with respect to the expected number of LOF variants under a neutral mutation model. In DeepLOF, we seek to estimate the distribution of *η_i_* from both genomic features and population genomic data. To this end, the DeepLOF model combines a feedforward neural network transforming genomic features and a likelihood function modeling the generation of LOF variants in human populations. Denoting x_*i*_ as the column vector of genomic features associated with gene *i*, the feedforward neural network describes the relationship between x_*i*_ and the prior distribution of *η_i_*. Denoting *y_i_* and *n_i_* as the observed and expected numbers of LOF variants in gene *i,* respectively, the likelihood function is defined as the probability of observing *y_i_* given *n_i_* and *η_i_*.

Specifically, we treat *η_i_* as a random variable ranging from 0 to 1 and utilize a beta distribution to describe its prior distribution,

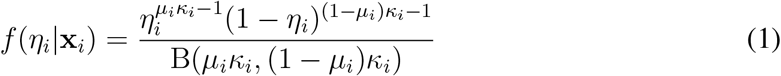

where *f* (*η_i_*|x_*i*_) is the probability density function of *η_i_* given feature vector x_*i*_; B is the beta function; *μ_i_* and *κ_i_* are the mean and concentration parameters of the beta distribution in gene *i*. It is worth noting that we employ an alternative parameterization of the beta distribution here^47^. The two shape parameters in the canonical parametrization of the beta distribution are equal to *μ_i_κ_i_* and (1 – *μ_i_*)*κ_i_*, respectively. Under the alternative parameterization, the mean of *η_i_* is equal to *μ_i_*, and the variance of *η_i_* decreases with increasing *κ_i_*. The alternative parametrization has been used in other Bayesian models due to the better interpretability of the mean and concentration parameters^47^.

In the feedforward neural network, we seek to model the relationship between x_*i*_ and the parameters of the beta prior distribution (*μ_i_* and *κ_i_*). There are two versions of feedforward neural network in DeepLOF: a nonlinear version with hidden layer and a linear version without hidden layer. Specifically, in the nonlinear version of DeepLOF, the hidden layers can be represented by the following equation,

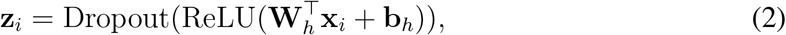

where z_*i*_ is the vector of hidden units; ReLU and Dropout are the the rectified linear layer48 and the dropout layer^49^; W_*h*_ and b_*h*_ are the weight matrix and the bias vector of the rectified linear layer. After the hidden layers, we add an additional layer to transform z_*i*_ into *μ_i_* and *κ_i_*,

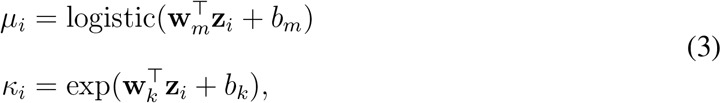

where w_*m*_ and *b_m_* are the weight vector and the bias term associated with *μ_i_*; w_*k*_ and *b_k_* are the weight vector and the bias term associated with *κ_i_*; the logistic function ensures that *μ_i_* ranges from 0 to 1; the exponential function ensures that *κ_i_* is positive.

In the alternative linear version of DeepLOF, the feedforward neural network does not include any hidden layer. Instead, we directly transform feature vector x_*i*_ into μ_*i*_ and *κ_i_*,

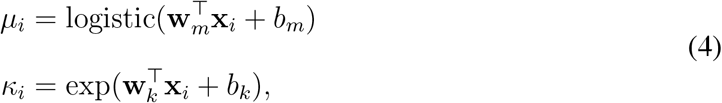

which is similar to equation 3 expect that z_*i*_ is replaced by x_*i*_. The linear DeepLOF model allows us to directly infer the associations of genomic features with LOF intolerance based on the negative values of weights in w_*m*_.

In the likelihood function, we seek to model the generation of LOF variants in human populations. Specifically, we assume that the observed number of LOF variants in gene *i* follows a Poisson distribution,

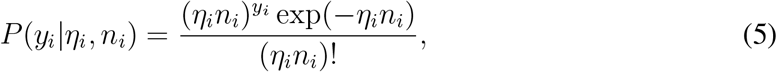

where *y_i_* and *n_i_* are the observed and expected numbers of LOF variants, respectively, and the mean of the Poisson distribution is equal to *η_i_n_i_*.

In the training step, the DeepLOF model combines the prior distribution (equation 1) and the likelihood function (equation 5) to obtain the marginal likelihood of the model,

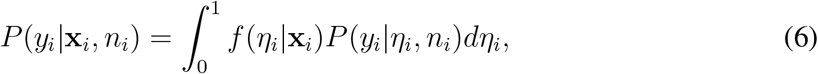

which represents the probability of observing *y_i_* LOF variants in gene *i* conditional on x_*i*_ and *n_i_*. It is worth noting that we omit the parameters of the feedforward neural network in this equation for the sake of notation simplicity. Because there is no analytical solution for the integral in this equation, we use the midpoint Riemann sum to approximately compute *P*(*y_i_*|x_*i*_,*n_i_*). To estimate the parameters of the feedforward neural network, we perform stochastic gradient descent on the following loss function,

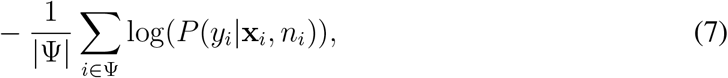

where Ψ and |Ψ| are the gene set and the number of genes in a mini-batch of data. We use the Adam algorithm50 for the mini-batch gradient descent and utilize early stopping and L2 regularization to avoid overfitting.

In the prediction step, we fix the parameters of the feedforward neural network to the optimal values from the training step. Then, we obtain the density function of the posterior distribution of *η_i_* using Bayes’ rule,

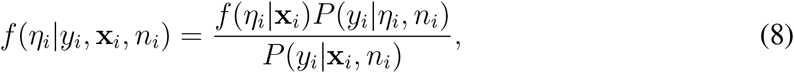

which represents our belief about *η_i_* after integrating genomic features and population genomic data. The mean of *η_i_* under the posterior distribution is equal to

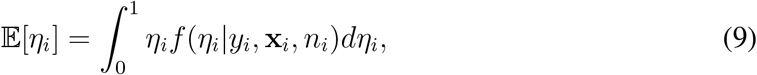

which we compute numerically using the midpoint Riemann sum. Finally, we define the DeepLOF score as 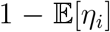. A higher DeepLOF score indicates that LOF mutations in the corresponding gene are subject to stronger negative selection.

### Genomic features

The training data of DeepLOF included 18 genomic features. First, we obtained five sets of epigenomic data from various cell types^20^. These data included ChIP-seq peaks of H3K9ac, H3K27me3, H3K4me3, and H2A.Z in promoter regions and promoter-enhancer interactions predicted by EpiTensor^51^. We defined H3K9ac, H3K27me3, H3K4me3, and H2A.Z signals as the average length of the corresponding ChIP-seq peak in a gene’s promoter across all cell types. We defined the enhancer number in a gene as the average number of promoter-enhancer interactions across all cell types. Second, we obtained four development-related gene categories from MSigDB25 (version 7.1). These gene categories included 1,029, 995, 508, and 1,131 genes from two GO categories24 (embryo development and central nervous system development) and two Reactome pathways33 (nervous system development and developmental biology). We converted each development-related gene category into a binary feature indicating whether each gene was included in the category. Third, we obtained a list of 1,254 transcription factor genes26 and a list of 3,431 genes encoding subunits of protein complexes^27^. We converted each gene list into a binary feature indicating whether each gene was included in the list. Fourth, we obtained promoter CpG density, promoter phastCons score, and exonic phastCons score from a previous study^15^. Fifth, we obtained mean gene expression level, tissue specificity (tau^34^), PPI degree28 from a recent study^52^. Finally, we obtained the UNEECON-G score from its original publication^32^.

We observed that several genomic features were nonnegative and had right-skewed distributions. Following a common practice in machine learning and statistics, we applied a log transformation to these features (Supplementary Table 1), 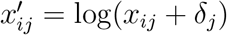, where *x_ij_* is the raw value of feature *j* in gene *i*, 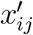 is the transformed feature, and *δ_j_* is the minimum observed positive value of feature *j*. Then, we standardized each continuous feature by subtracting its mean and dividing by its standard deviation. We imputed missing values of each feature with the mean of the non-missing values.

### Model training

We obtained the observed number of LOF variants in each protein-coding gene and the expected number under a neutral mutation model from gnomAD (version 2.1.1). We combined the expected and observed numbers of LOF variants with the 18 genomic features to build a dataset of 19,197 genes for model training. We randomly split these genes into a training set (80% genes) and a validation set (20% genes). We trained the DeepLOF model on the training set and used a grid search to tune hyperparameters in the validation set. In the training of the linear DeepLOF model, these hyperparameters included the L2 penalty (0, 10^-2^, 10^-3^, 10^-4^, 10^-5^, 10^-6^) and the learning rate of the Adam algorithm (10^-3^, 10^-4^, 10^-5^). In the training of the nonlinear DeepLOF model, we added an additional hyperparameter, *i.e.,* the number of hidden units in the feedforward neural network (64, 128, 256, 512, 1,024). We fixed the dropout rate to 0.5. We computed the contribution scores of genomic features using the optimal linear DeepLOF model with the lowest loss in the validation set. We computed the DeepLOF score using the nonlinear model with the lowest loss in the validation set. The optimal nonlinear model had a lower loss than the optimal linear model.

### Comparison with other methods in predicting disease genes

We evaluated the performance of DeepLOF and eight alternative methods, including LOEUF^5^, pLI^2^, mis-z^1^, RVIS^36^, GeVIR^6^, CoNeS^7^, VIRLOF^6^, and UNEECON-^G32^, in predicting essential genes and dominant-negative genes. We obtained LOEUF, pLI, and mis-z scores from the gnomAD database5 (version 2.1.1). We obtained the RVIS score trained on the ExAC dataset2 from dbNSFP53 (version 4.0). We obtained the other gene scores from the corresponding publications.

We obtained 311 ClinGen haploinsufficient genes and 404 mouse genes where heterozygous knockouts resulted in lethality from the GitHub repository for gnomAD (https://github.com/macarthur-lab/gnomad_lof/). Then, we obtained 18,797 human-mouse orthologs from the Mouse Genome Database^22,54^ and used these data to map the mouse essential genes to the human genome, resulting in 397 human orthologs of mouse essential genes. We obtained 683 human genes deemed essential in cell lines and 913 genes without significant fitness effects in cell lines from the GitHub repository for the MacArthur Lab (https://github.com/macarthur-lab/gene_lists). We obtained 364 OMIM dominant-negative genes from a previous study^36^.

For each of these positive gene sets, we constructed a negative gene set with matching size. Specifically, for the 311 ClinGen haploinsufficient genes, the 397 human orthologs of mouse essential genes, and the 364 dominant-negative genes, we considered all other human genes to be negative. For the 683 human genes deemed essential in cell lines, we considered the 913 human genes without significant fitness effects in cell lines to be negative. Then, we used MatchIt35 to match each positive gene with a negative gene of similar expected number of LOF variants. Finally, we used ROCR to plot the receiver operating characteristic curves and calculate the AUCs for all computational methods in the matched gene sets^55^. We evaluated the statistical significance of the difference in AUC using the DeLong test^56^.

### Evaluation of the 109 LOF-tolerant genes uniquely predicted by DeepLOF

We obtained comparable numbers of LOF-intolerant genes from DeepLOF, LOEUF, VIRLOP, and CoNeS. First, we obtained 2,835 LOF-intolerant genes from LOEUF using an established cutoff of 0.35 (ref. 5). To obtain similar numbers of LOF-intolerant genes from the other methods, we used cutoffs of 0.835, and −1.11, and 15 for DeepLOF, CoNeS, and VIRLOF percentile scores, respectively. Given these cutoffs, DeepLOF, CoNeS, and VIRLOF predicted 2,817, 2,847, and 2,817 LOF-intolerant genes, respectively. We retained LOF-intolerant genes with ≤ 10 expected LOF variants for downstream analysis.

We evaluated the enrichment of ClinGen haploinsufficient genes, human orthologs of mouse essential genes, and human genes essential for the survival of cell lines in the 109 LOF-intolerant genes uniquely predicted by DeepLOF. For each essential gene set, we defined the other genes as nonessential genes. Also, we defined LOF-tolerant genes as those genes with ≤ 10 expected LOF variants and not predicted to be LOF-intolerant by any method. We evaluated the enrichment of each essential gene set in the 109 LOF-intolerant genes using the log odds ratio, 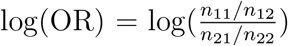, where *n*_11_, *n*_12_, *n*_21_, and *n*_22_ are the numbers of essential genes predicted to be LOF-intolerant, nonessential genes predicted to be LOF-intolerant, essential genes predicted to be LOF-tolerant, and nonessential genes predicted to be LOF-tolerant, respectively. We defined the confidence interval of the log odds ratio as log(OR) ± 1.96 × SE, where SE is the standard error of the log odds ratio and is equal to 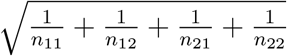.

We evaluated the depletion of the 109 LOF-intolerant genes uniquely predicted by DeepLOF in benign genomic deletions. To this end, we obtained clinical structural variants from the nstd102 study in dbVar39 and retrained 5,649 benign deletions overlapping coding regions of genes from GENCODE57 (version 19). Then, we computed the proportion of benign deletions overlapping at least one of the 109 LOF-intolerant genes. To examine whether the proportion of overlapping deletions was smaller than the expectation under a null model that the 109 LOF-intolerant genes are nonessential. We performed a permutation test with 10,000 permutations. In each permutation, we randomly selected 109 genes with ≤ 10 LOF variants and computed the proportion of deletions overlapping with the random genes. The one-tailed *P*-value of the permutation test was defined as the fraction of permutations where the proportion of deletions overlapping random genes was equal to or smaller than the observed proportion in empirical data.

## Data availability

The training data, DeepLOF score, and predicted LOF-intolerant genes with ≤ 10 LOF variants from this study are available as Supplementary Data files.

## Code availability

The DeepLOF model and companion documentation are available at https://github.com/yifei-lab/DeepLOF.

## Acknowledgments

Research reported in this publication was supported by the National Institute of General Medical Sciences of the National Institutes of Health under Award Number R35GM142560 and by the Pennsylvania State University. The content is solely the responsibility of the authors and does not necessarily represent the official views of the National Institutes of Health.

