## Supplementary Figures and Tables for "A deep learning framework for predicting human essential genes from population and functional genomic data"

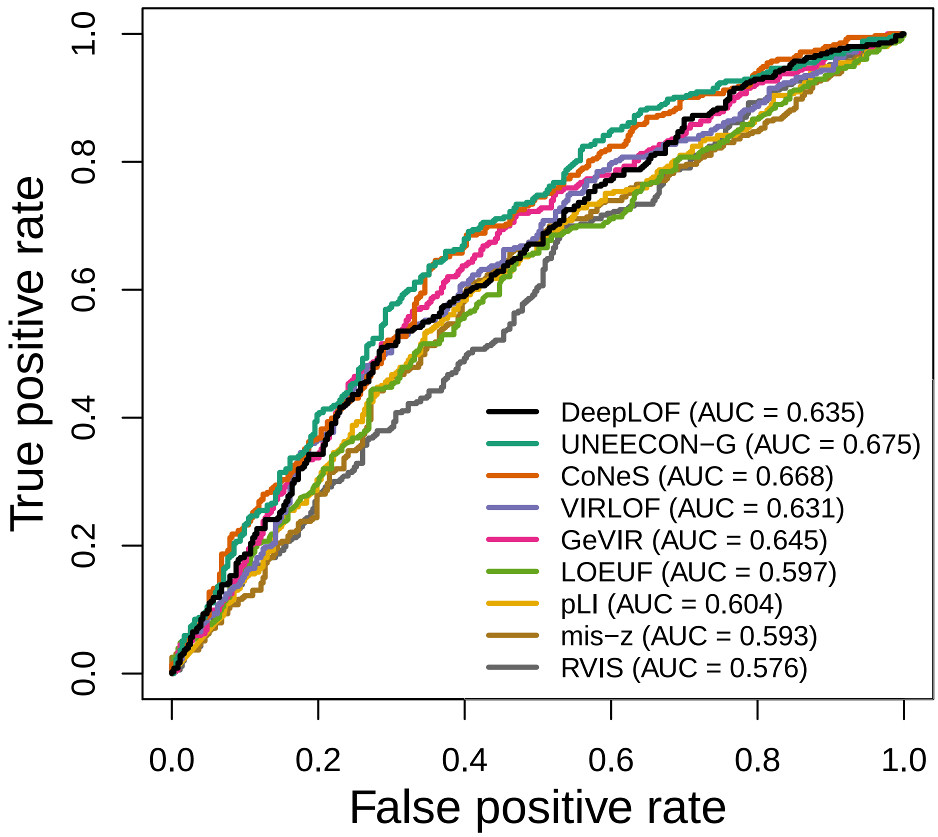


Supplementary Fig. 1: Performance of DeepLOF and alternative methods in predicting dominant negative genes.

Supplementary Table 1: Genomic features for model training.

| Feature name | Feature type | Log transformation | Data source |
|---|---|---|---|
| H3K9ac signal | Continuous | Yes | [1] |
| H3K27me3 signal | Continuous | Yes | [1] |
| H3K4me3 signal | Continuous | Yes | [1] |
| H2A.Z signal | Continuous | Yes | [1] |
| Enhancer number | Continuous | Yes | [1] |
| GO embryo development | Binary | N/A | [2] |
| GO central nervous development | Binary | N/A | [2] |
| Reactome nervous system development | Binary | N/A | [2] |
| Reactome developmental biology | Binary | N/A | [2] |
| Transcription factor | Binary | N/A | [3] |
| Protein complex | Binary | N/A | [4] |
| Promoter CpG density | Continuous | No | [5] |
| Promoter phastCons score | Continuous | No | [5] |
| Exonic phastCons score | Continuous | No | [5] |
| Mean expression level | Continuous | Yes | [6] |
| Tissue specificity (tau) | Continuous | No | [6] |
| PPI degree | Continuous | Yes | [6] |
| UNEECON-G | Continuous | No | [7] |

Supplementary Table 2: Statistical significance of the differences in AUC between DeepLOF and alternative methods in predicting essential genes. The numbers represent $P$-values from the DeLong test. *: $P < 0.05$; **: $P < 0.01$; ***: $P < 0.001$.

| DeepLOF | ClinGen haploinsufficient genes | Human orthologs of mouse essential genes | Human essential genes in cell lines |
|---|---|---|---|
| *vs.* UNEECON-G | 1.332e-05 *** | 5.835e-04 *** | 5.577e-10 *** |
| *vs.* CoNeS | 8.679e-04 *** | 3.688e-03 ** | 4.068e-21 *** |
| *vs.* VIRLOF | 6.909e-07 *** | 4.766e-06 *** | 1.260e-02 * |
| *vs.* GeVIR | 1.965e-08 *** | 2.181e-05 *** | 7.913e-05 *** |
| *vs.* LOEUF | 6.601e-10 *** | 1.073e-09 *** | 3.151e-12 *** |
| *vs.* pLI | 1.090e-09 *** | 3.263e-10 *** | 6.640e-50 *** |
| *vs.* mis-z | 8.264e-22 *** | 1.435e-14 *** | 5.962e-09 *** |
| *vs.* RVIS | 8.893e-23 *** | 1.834e-16 *** | 2.067e-15 *** |
